# Not Extremely Plastic: Testing the Limits of Morphological Plasticity in Fungal Mycelia in Response to Soil Grazers

**DOI:** 10.1101/2025.01.14.632904

**Authors:** C.A. Aguilar-Trigueros, L. Boddy, M.D. Fricker

## Abstract

Phenotypic plasticity is widespread among living organisms, but exceptionally high levels of plasticity are rarely observed in nature. While plasticity can enhance fitness in fluctuating environments, it comes with substantial costs that limit both its expression and evolution. Most of what we know about these limits comes from studies on animals and plants, which typically have fixed body plans. In contrast, modular organisms such as filamentous fungi possess an undetermined, decentralized morphology that allows individuals to adjust their phenotype in response to local environmental conditions. This flexibility has led to speculation that fungi exhibit "extreme phenotypic plasticity," making them an ideal system for studying plasticity mechanisms—yet empirical tests of this assumption remain scarce. To address this gap, we analyzed grazer-induced morphological plasticity in four cord-forming fungal species exposed to soil microfauna with differing feeding behaviors, using a phenotypic trajectory analysis framework and tested alternative hypotheses: One hypothesis is that the flexibility of fungal network development would allow species to converge on a common "grazing-resistant-phenotype". Alternatively, the developmental plasticity for each species might be heavily constrained within species-specific limits. Our results showed that fungal species accounted for most of the total morphological variation, while grazer induced morphologies remained relatively aligned to non-grazed morphological trajectories. Instead of convergence, we found that grazed-phenotypes were more dissimilar across species, reflecting distinct developmental trajectories within species-specific limits. The extent of plasticity varied widely with a maximum of 30% of the morphological variation attributable to grazing pressure. Nevertheless, these findings suggest that morphological plasticity plays a relatively minor role in response to grazing for most species. The dominant species signature and constrained plasticity supports the use of morphological attributes of mycelia as functional traits in ecological studies. Our findings provide new evidence on the extent of plasticity of modular organisms and highlight the need to extend plasticity theories to include organisms with non-determinate, modular body types like fungi.

## Introduction

Determining which organisms achieve high levels of phenotypic plasticity has long been a central question in ecology and evolution (Sultan 2003, 2021; West-Eberhard 1989). In fluctuating environments, plasticity—the ability of a single genotype to alter its phenotype in response to environmental change (Pigliucci & Müller 2010)—is crucial for maintaining or enhancing fitness in the new conditions (Forsman 2015; Pfennig 2021). While all living organisms exhibit some degree of plasticity, extreme or "perfect" forms of plasticity are rarely found in nature (Gomulkiewicz & Stinchcombe 2022; Murren *et al*. 2015). This limited occurrence is thought to reflect the inherent costs of plasticity, including the energetic burden of modifying phenotypes and maintaining the regulatory systems needed to sense and respond to environmental variation (Auld *et al*. 2010; Schneider 2022). Yet, theory suggests that modular organisms—those lacking a fixed adult body plan and instead grow by iteration of repeated units or modules—may be less constrained in their plasticity compared to organisms determined body plans like animals (Murren *et al*. 2015). This modularity allows these organisms to adjust development and structure more flexibly in space and time compared to unitary organisms, potentially allowing them to evolve much higher degrees of plasticity. Yet, modular lineages remain underrepresented in studies of plasticity, leaving it unclear whether and how often extreme plasticity evolves in such systems.

Filamentous fungi represent a prime group in which to explore the potential for extreme phenotypic plasticity in modular organisms. This group is a dominant lineage in terrestrial ecosystems (Anthony *et al*. 2023) but have a fundamentally different body plan from that of animals as they grow as networks of interconnected filaments (hyphae) that together form a mycelium (Fricker *et al*. 2017a) without a determined body morphology. Empirical evidence shows that fungi can adjust their morphology in response to environmental variation. In extreme cases, fungi can switch between unicellular (yeast-like) and filamentous (mycelium-like) forms depending on conditions such as temperature, pH, or nutrient availability (Boyce & Andrianopoulos 2015; Gauthier 2015). Within filamentous phenotypes, shifts in space occupancy, colony shape, and pigmentation have been observed in response to resource heterogeneity and environmental stress (Boddy 1999; Slepecky & Starmer 2009; Veresoglou *et al*. 2018). This morphological flexibility has led to the proposition that fungi exhibit “extreme phenotypic plasticity”(Klein & Paschke 2004; Slepecky & Starmer 2009; Ugalde & Rodriguez-Urra 2014), and that because of such high levels of plasticity fungi are “the most adaptable phylogenetic lineage in the Tree of Life” (Coleine *et al*. 2022). Yet research quantifying the extent of plasticity in this group has remained scarce (Alster *et al*. 2021; Behm & Kiers 2014).

While plasticity can be evaluated in response to any environmental cue, the universe of potential environmental variation leads to a factorial explosion in experimental design. Particularly for fungi, environmental variation includes a plethora of microclimatic conditions, resource heterogeneity and biotic interactions. Variation in resources alone stems from differences in resource type, quantity, accessibility, quality and spatial distribution. However, certain biotic response, such as predator-induced plasticity—where a genotype alters its phenotype in response to the presence of a predator—provides a more tractable experimental system where the extent of plasticity can be probed using a simpler presence-absence experimental design. This approach has been widely used with animals to develop model systems to understand the ecology and evolution of plasticity (Hoverman & Relyea 2007; Tollrian & Harvell 1999). These studies typically involve quantitative comparison between the phenotype of clonal genotypes in the presence and absence of a predator. Classic examples with the model animal *Daphnia* show that exposure to predators induce pronounced changes in its morphological traits (Spitze & Sadler 1996), a pattern that is observed across several other animals including rotifers (Gilbert 2013; Yin *et al*. 2017), dragonflies (Flenner *et al*. 2009), snails (Auld & Relyea 2011) and aphids (Sentis *et al*. 2019). Likewise in plants there is a growing number of studies investigating how the presence of herbivores triggers morphological plasticity (Dorey & Schiestl 2022; Fernández De Bobadilla *et al*. 2022). Furthermore, the changes induced depend on the feeding habits of the herbivore (Lebbink *et al*. 2024).

Fungal grazing by soil fauna is analogous to animal predator-prey or plant-herbivore interactions, thus making it a candidate to study plasticity of fungal phenotypes. In a similar manner to experiments conducted in animals, fungal plasticity can be assessed by growing clonal replicates of the same fungal genotype in the presence or absence of naturally occurring grazers, using densities that reflect natural conditions (Crowther *et al*. 2012). There is also evidence that two basic elements for plastic morphological response of fungal morphology to grazing are present in fungi. First, grazing reduces fungal fitness (Tordoff *et al*. 2006, 2008) thus, has the potential to impose evolutionary pressure for the emergence and maintenance of grazer-induced plasticity. Such pressures are imposed by grazing by various different animals, including nematodes, collembola, millipedes and isopods. As each of those grazers have different feeding behaviour, plastic responses may be specific to the behavior of these organisms (Crowther *et al*. 2012).

Second, there is evidence that fungi can detect the presence of grazers. For example, (Schmieder *et al*. 2019) identified sensory pathways activated in the model mushroom *Coprinopsis cinerea* when exposed to a fungivorous nematode. Similarly, the edible oyster mushroom (*Pleurotus ostreatus*) not only detects the presence of nematodes but also releases toxins and develops traps to immobilize and consume them under low-nitrogen conditions (Lee *et al*. 2020).

Until recently one of the main challenges of studying morphological plasticity of fungi has been the difficulty of quantitatively describing the structure of its modular body, that is, their mycelial networks. However, semi-automated pipelines have now been developed that provide multiple metrics describing fungal network morphology and predicted function (Dikec *et al*. 2020; Du *et al*. 2016; Fricker *et al*. 2017b; Heaton *et al*. 2012b; Sten *et al*. 2024; Vidal-Diez De Ulzurrun *et al*. 2015), including mycelia foraging for new resources on the surface of soil microcosms which represent a more realistic environment to measure behavioural responses than homogenous high resource growth media (Aguilar-Trigueros *et al*. 2022). Network-forming fungi do not forage by active movement in the way that animals do, but rather they explore their surrounding environment by growing out of a resource patch even when the surrounding area has no immediate resources available (Bielčik *et al*. 2019). An extreme case of this type of foraging ability is observed for “cord-forming fungi” that form hyphal aggregates termed cords which can be several mm thick and cm to meters long (Boddy 1999), forming macroscopically visible systems often covering large areas in the field (Fricker et al., 2007; Yafetto, 2018).

Nutrients and water are transported within these cords between research patches and to and from mycelial fronts (Tlalka *et al*. 2002). Thus these group of fungi provide a tractable model system to investigate morphological plasticity, particularly to grazing, given the numerous studies reporting the impact of grazing on their fitness (reviewed by (Crowther *et al*. 2012). Possible plastic morphological responses of cord-forming mycelia to grazing include the formation of new interconnections providing alternative routes for transport, development of a denser mycelium and thickening of cords to limit damage (Rotheray *et al*. 2008).

To determine the extent of grazer-induced plasticity in these fungi, we reanalyzed an existing image dataset of cord-forming mycelia from four fungal species exposed to grazing by a range soil micro-fauna with different feeding behaviour (Crowther *et al*. 2011). The original study quantified changes in radial extension rate, hyphal coverage, fractal dimension, and wood decay rate and detected significant fungal-grazer interactions (Crowther *et al*. 2011). The experimental design parallels reaction norm studies typically used to evaluate predator-induced plasticity (i.e. clones of the same genotype growing with or without grazers) and allowed us to quantify changes in the network architecture of each fungal genotype in response to grazers. Specifically, we measure a suite of mycelial morphological traits that capture the network architecture of the fungal mycelium from time series images (Aguilar-Trigueros et al., 2022; Fricker et al., 2007; Fricker et al., 2017). We used phenotypic trajectory analysis to quantify the extent of developmental plasticity in response to grazing. This method, widely used to study both evolutionary (Adams & Collyer 2009) and developmental plasticity (Metcalfe 2024) using multiple traits (Hoverman & Relyea 2007), has been suggested as a powerful tool for studying fungal plasticity (Behm & Kiers 2014).

Given the high degree of morphological plasticity expect in fungal mycelia and the strong fitness pressures imposed by grazing, we tested two alternative hypotheses about how these modular organisms express plasticity in response to environmental stress (Fig. 1). In the first scenario—extreme plasticity—we hypothesized that different fungal species exposed to the same grazing pressure would converge on a common morphological trajectory, producing a shared, potentially optimal “grazing-resistant” phenotype. Under this scenario, plastic responses effectively erase the original phenotypic distinctions between taxa and set their developments into a new path. In contrast, under a constrained plasticity scenario, we expected each species to modify its morphology in response to grazing but remain close to its species-specific developmental trajectory. This scenario would reflect intrinsic limits to plasticity shaped by lineage-specific constraints highlighting evolutionary fixed traits. By disentangling these alternatives, our goal is not only to assess the extent of plasticity in filamentous fungi, but also to uncover the extent that modular growth forms mediate adaptive responses to environmental change.

**Figure 1.**
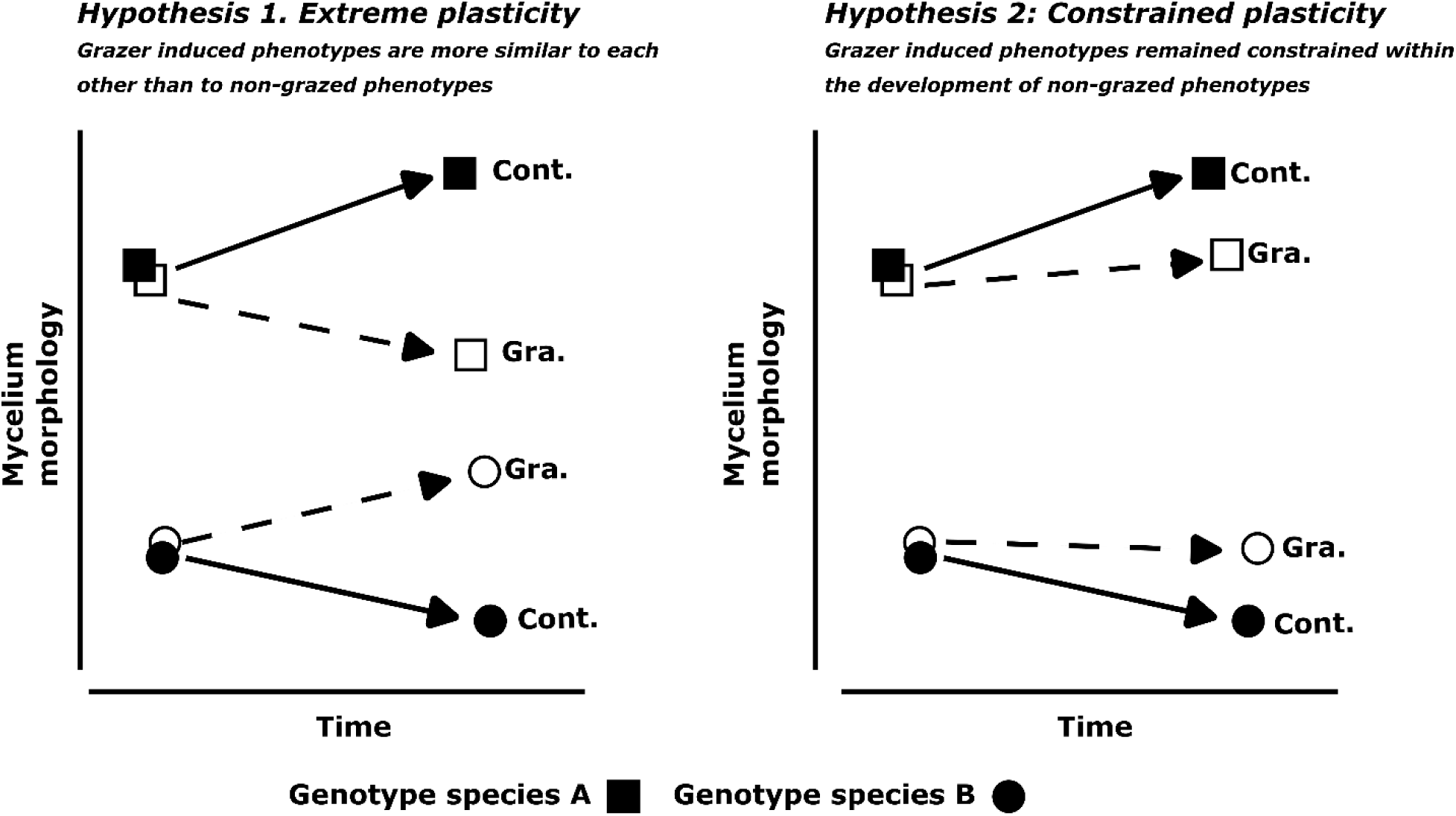
Conceptual representation of hypotheses predicting fungal morphological plasticity in response to grazing. Under Hypothesis 1 (Extreme plasticity), grazer-induced phenotypes (open symbols, dotted lines) diverge so strongly from their non-grazed counterparts (closed symbols, solid lines) that they resemble grazer-induced phenotypes of other genotypes more than their own non-grazed phenotype. Under Hypothesis 2 (Constrained plasticity), grazer-induced phenotypes still differ from non-grazed ones, but remain within the trajectory defined by their genotype’s inherent morphological trajectory.

## Materials and methods

### Experimental setup

The data for this study were derived from a time-series experiment initially designed to investigate fungal grazing interactions (Crowther *et al*., 2011). The data set includes isolates from four species of wood-decomposing fungi: *Phallus impudicus*, *Phanerochaete velutina*, *Hypholoma fasiculare*, and *Resinicium bicolor* that were originally isolated from woodlands in the southern UK and are maintained in the Cardiff culture collection (Boddy 1999). Mycelium growth occurs by extension of cords (in length) and by cord thickening. In addition, new cords are created through branching, while some cords fuse with one another to cross-link the network. The resources necessary to sustain growth and metabolism are transported out of the wood block, which provides the only carbon-resource in the microcosm. Differences in morphology arise by changing the rates of cord extension, thickening, branching and fusion. Briefly, for each species, 30 clones from the same genotypes were cultivated from 20 × 20 × 10 mm beech blocks and allowed to grow across compressed soils in lidded 245 mm × 245 mm trays at 18C. The mycelia grew for 8-12 days until at least half of the replicates had reached a diameter of 160-200 mm (see supplementary material for a detailed description of the imaging settings). Five clones per species (i.e. replicates) were then exposed to five types of grazing pressure, along with a set that was maintained grazer free that served as control. Grazing pressure (i.e. grazing treatment) consisted of five grazing levels: collembola (*Folsomia candida*), millipedes (*Blaniulus guttulatus*), woodlice 1 (*Oniscus asellus*), woodlice 2 (*Porcellio scaber*), nematodes (*Panagrellus redivivus*) and a grazer-free control.

To monitor mycelial development over time, images were captured at the start of the experiment (day 1, before grazer addition), and subsequently on days 2, 4, 8, 16, and 32. Here we restrict analysis up to the 8-day time-point to ensure the network architecture was not perturbed by physical constraints as some species reached the edge of the petri dish beyond this day. Images were captured with pixel sizes ranging from 80 to 155 µm.

### Quantitation of mycelial network architecture

We use a set of image processing algorithms to extract the mycelial morphology/architecture from pictures as described in (Aguilar-Trigueros *et al*. 2022). Briefly, each picture was loaded into a graphical user interface (GUI, Fig. S1) and processed through a standard pipeline that included background correction, network enhancement, segmentation, skeletonization, width estimation and network graph representation using protocols optimized across a range of biological network systems (Fricker, et al., 2017; Obara et al., 2012; Pain et al., 2019; Xu et al., 2021). For these macroscopic networks with cords of different thickness and reflectivity, the mean-phase angle from intensity-independent phase congruency analysis gives marginally better network enhancement performance compared to other standard approaches such as second-order anisotropic Gaussian (Lopez-Molina *et al*. 2015; Vidal-Diez De Ulzurrun *et al*. 2015) or Bowler-Hat (Oyarte Galvez *et al*. 2025) algorithms used for networks grown on transparent media, although all are implemented in the software. The software package and complete manual explaining in detail the steps involved is freely available from Zenodo (DOI: <u>10.5281/zenodo.5187932</u>). The output of this pipeline is a network graph representation of the mycelia where the tips, cord branching and cord fusion events are as nodes whilst the cords themselves form ‘edges’ connecting the nodes. (Fig. S1). The length and width of each edge was automatically measured, whilst the (*x,y*) position and branching angle were recorded for each node. It was not possible to resolve edges within the inoculum, so these are represented as strong connections between each incident edge on the resource and a central node (termed the ‘Root’ of the network). This information was exported in two tables as .csv and .xlsx files containing the cord (edge) information and the tips, branching and fusion points (node) information. The node and edge tables from the image processing pipeline were imported into R (using the *read_excel* function from the readxl package (Wickham *et al*. 2019) and used to construct a spatially explicit transport network embedded in 2D space given by the Euclidean coordinates of the nodes using the igraph package in R (Csardi G & Nepusz T 2006) (Figure 2. See also details in the code available in the github repo https://github.com/aguilart/Fungal_Networks).

**Figure 2.**
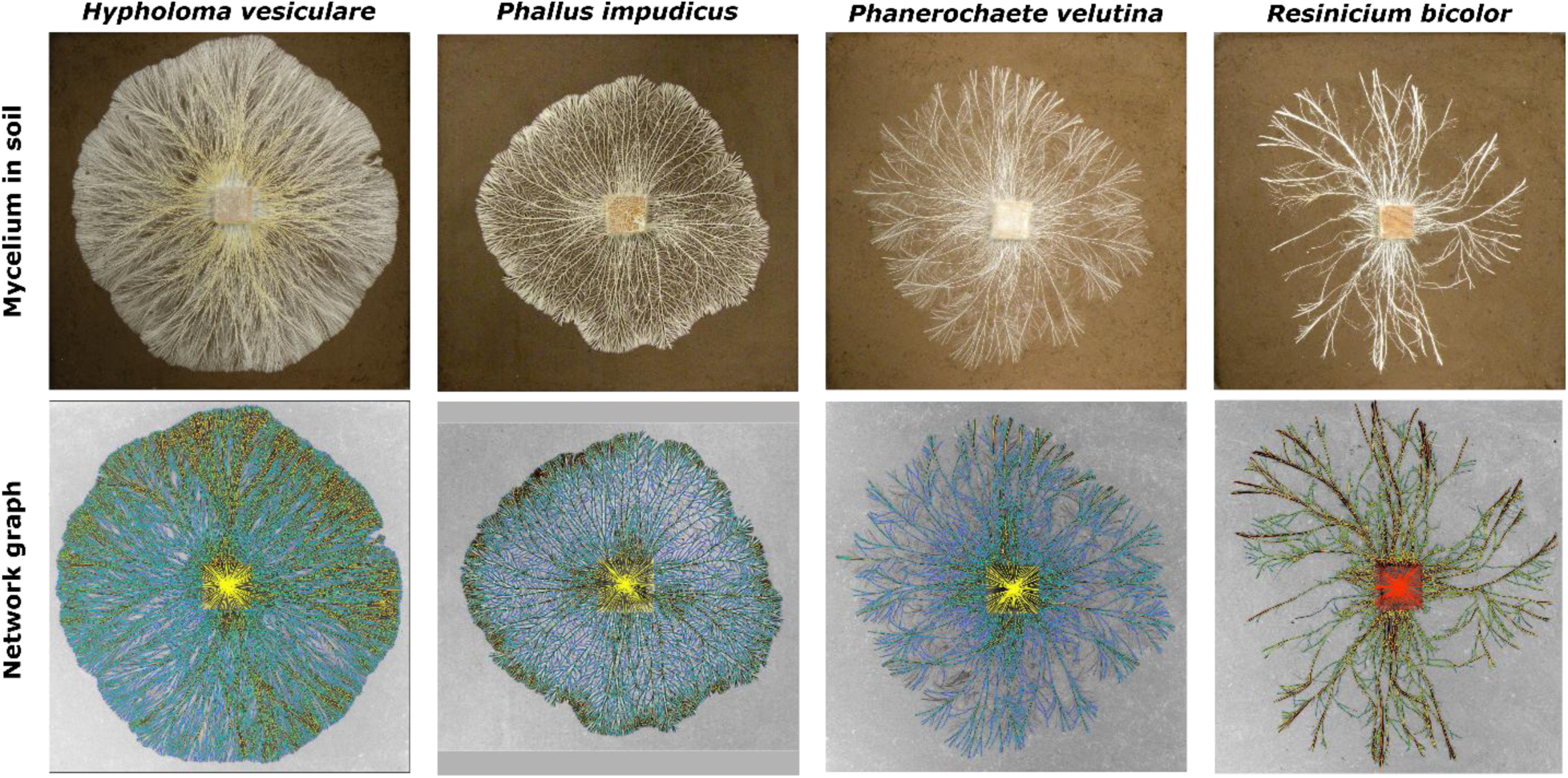
Translating cord-forming fungi into a weighted transport network. *Top row*: Photographs of mycelia from the four fungal species used in this study. The fungi grow out from 2×2×1 cm wood blocks, forming millimeter-thick cords that explore the surrounding environment in search of new resources. Fungi were grown under laboratory conditions on compressed soil in 245 mm × 245 mm trays. *Bottom row*: Network graphs overlaid on inverted grayscale images of the mycelia. These graphs were generated following intensity-independent phase congruency edge enhancement, local adaptive thresholding and skeletonization of the cords and integrated granulometry to measure the cord width. Cords are represented as network edges connected by nodes comprising tips, branching points, and fusion points (anastomoses). Cord width is visualized with color: green for thick cords, blue for thin cords. For visualization purposes, the inoculum is represented as multiple linear edges in yellow, but these are not included in the analysis.

### Traits used to define mycelial network morphology

To measure mycelium morphology, we first identified the cords that represent the main routes of the network as the minimum spanning tree (MST) of cords that confer the least transport resistance from the wood block to the foraging tips at the front. Cords that were not part of the MST were recorded as secondary routes that enhance cross-linking between the main cord routes. We measured a set of 22 network metrics that reflect the overall colony morphology, network heterogeneity, space filling and connectivity, predicted transport and predicted robustness to damage (Table 1, S1). Predicted transport in the network was simplified as unidirectional fluid flow from the wood block to the growing front of the network by assuming cords can be represented as a contiguous network of interconnected bundles of cylindrical shape. The predicted resistance to flow scales with cord length and inversely to the square of the radius of the cord (i.e. *resistance µ l/r^2^*) {Bebber, 2007 #1793;Fricker, 2007 #2029}. Thus, long and thin cords provide higher resistance. This is a simplification of transport dynamics of mycelia but matches well with empirical distributions of radiolabeled nutrient movement in these networks (Heaton *et al*. 2010, 2012a).

**Table 1.**
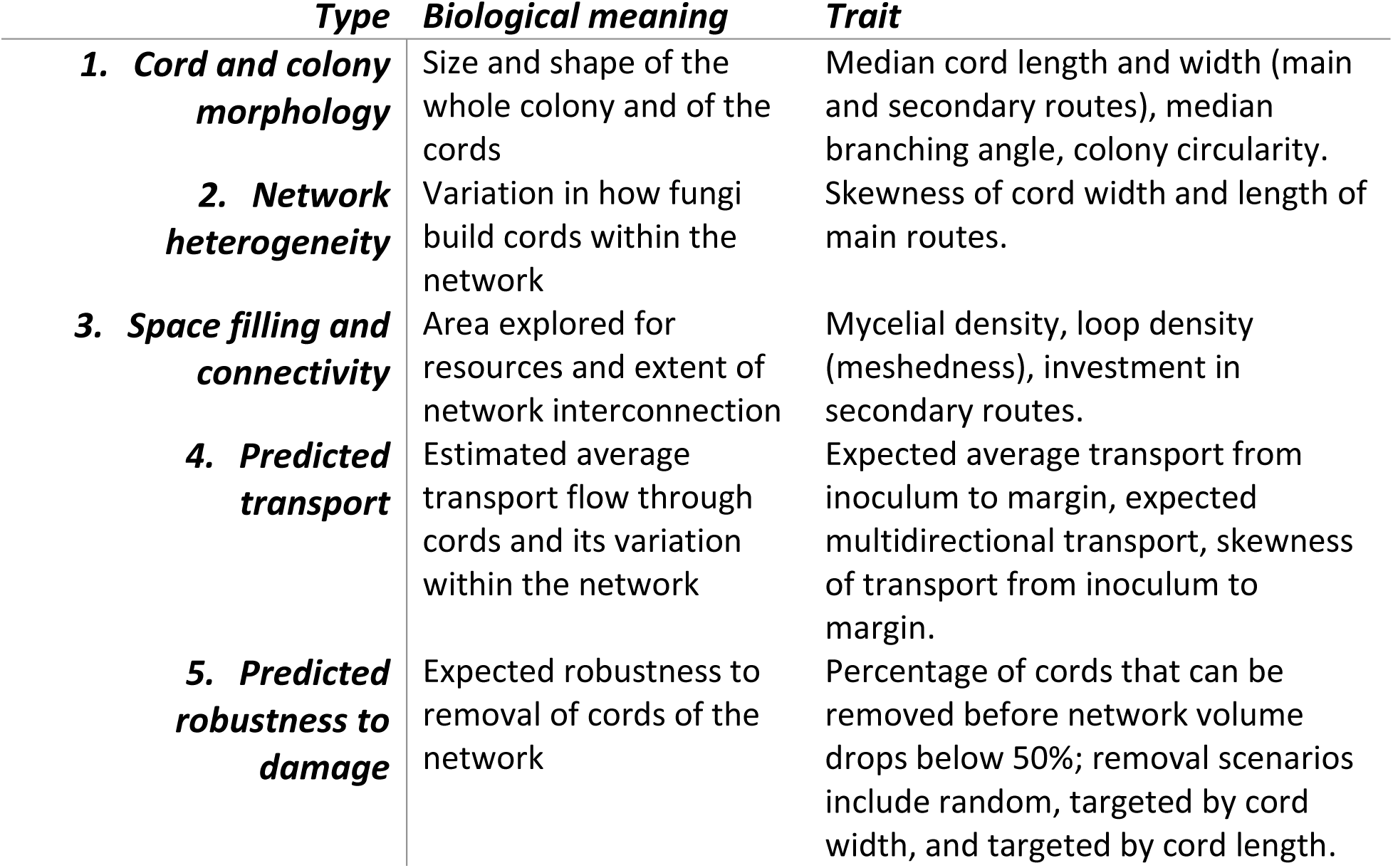
Description of the five trait types used in this study. . These traits summarize different patterns of the whole mycelium (see table S1 for full list of 21 metrics measured).

### Phenotypic trajectory analysis to measure morphological developmental plasticity

Phenotypic trajectory analysis relies on the use of multivariate ordination methods to summarize the change through time in phenotypes across multiple traits combined using ANOVA-like frameworks for statistical testing (Adams & Collyer 2009; Behm & Kiers 2014). In this study, we used three types of permutational multivariate analyses of variance models (PERMANOVA) to determine level of grazer-induced plasticity across the fungal species using the 22 metrics. In all the models, we controlled for initial differences in inoculum growth at the start of the experiment by using the first two axes from a principal component analysis (PCA) of traits at time 1 (i.e. immediately before the addition of the grazers) as covariate in the PERMANOVA.

In the first and more general model, we tested whether induced changes in the development of the morphology (plasticity) converge to a common grazing-resistant -phenotype across all four species (Hypothesis 1, extreme plasticity) or whether plastic response remained constrained close to the trajectories of non-grazed phenotype per species. In this model, time, fungal species identity, and grazer type were used as fixed factors and PC1 and PC2 of the traits used at the start of the experiment to control for initial conditions (see table 2 for the full formula used in this model). If fungal species converge to a common phenotype, we expect to detect a strong (and significant) main effect of grazer together with an interaction between grazer and time. That is, strong interaction grazer x time suggests more deviation of the development of the morphology of grazer-exposed clones compared to the grazer-free clones, while strong main grazer effect suggest similarity of morphology of the grazed phenotype while.

In the second model, we determine which grazers hand the strongest effect in inducing a common phenotype at the end of the experiment. We ran five PERMANOA models (i.e. one for each grazer) where the morphological traits of all four fungal species were used as response variables while the only fixed factor was the grazer treatment with two levels (presence or absence). Convergence would be shown by a significant, large effect size of grazer identity. To further visualize and validate the output of these models, we complemented this model by testing whether the grazers increase the similarities among fungal species compared to the control. This test was done by measuring the distances between each fungal species across the respective grazer treatments to their multivariate mean (centroids) followed by a permutation based-test for multivariate homogeneity of group dispersions (implemented with the function *betadisper* from the package *vegan* in R). If grazers increase the similarity (or decrease it) it would be shown as a strong and significant reduction in the distances between each species and their respective centroids.

The third model tested which fungal species had the most plastic response to grazers. We ran PERMANOVA models for each fungal species. Thus, this test measured the extent of development plasticity of fungal species irrespective convergence to a common phenotype. Finally, for the fungal species showing the largest plastic responses, we ran a set of models with each grazer separately to identify which trait drove the plastic responses.

In all PERMANOVA models, we used a forward model selection approach to determine which fixed factors should be retained. We used Principal Component Analysis (PCA) to visualize the changes in the phenotype through time. Analysis of the distances to each centroid were done using Euclidean distances. Effect sizes were measured as the proportion of explained variance of the fixed factors. PERMANOVA, PCA, analysis of distances to centroids, model selection and effect sizes were calculated using functions from the vegan package (Oksanen *et al*. 2007). All statistical analysis were conducted in R (R Core Team 2013).

## Results

### Support for constrained plastic response rather than extreme plasticity

Species identity had the strongest effect, explaining up to 90% of the variation in morphology across all treatments and time points using model 1 (Fig. 3). We found significant effects of grazer identity and an interaction with time, but neither effect was larger than 5% of the total variation (Table 2).

**Figure 3.**
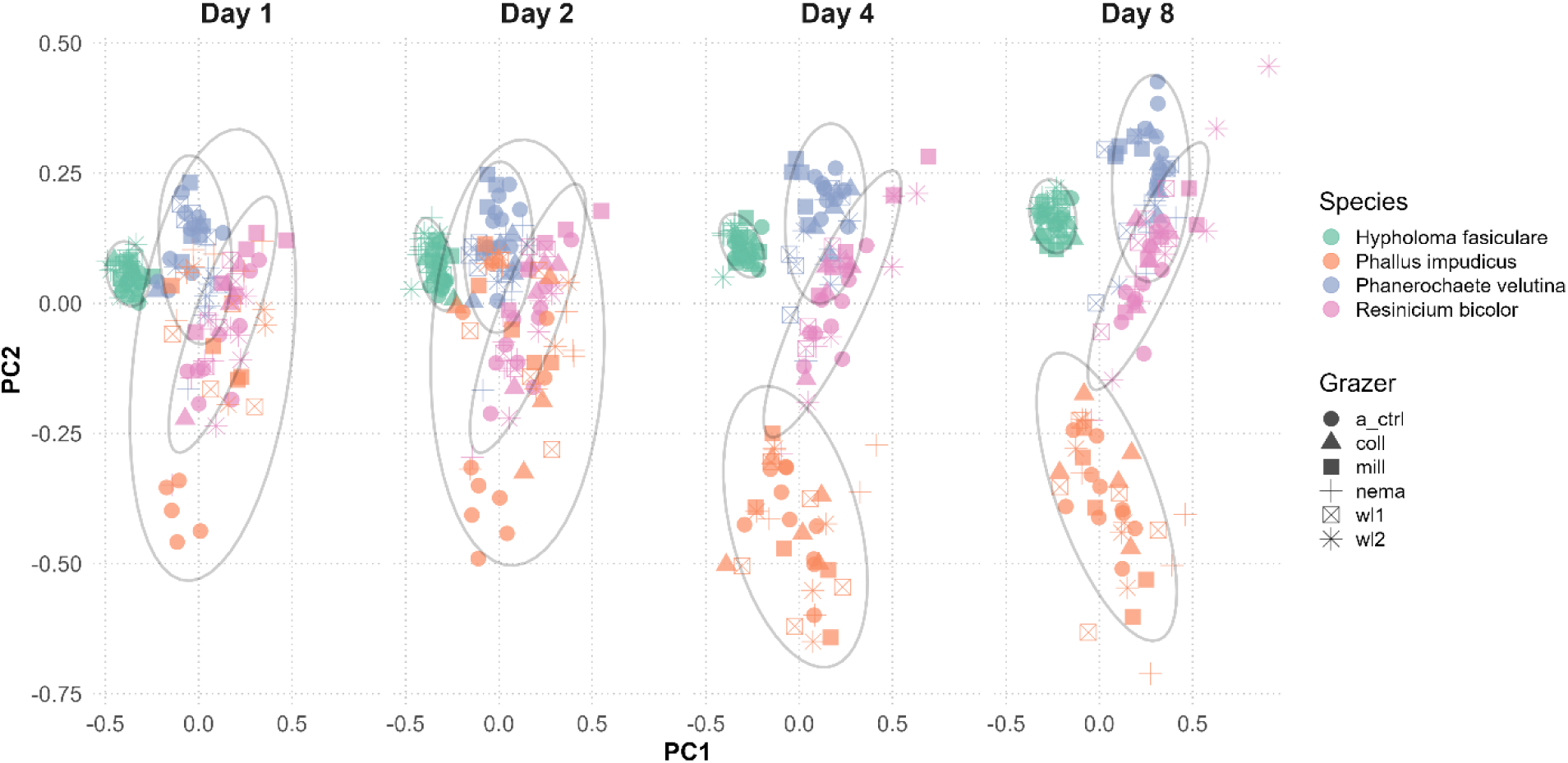
Figure 3. PCA showing the temporal development of mycelial morphology in four fungal species. At the start of the experiment, species show similar mycelial traits, but as time progresses, distinct species-(genotype-) specific morphologies emerge. While the presence of grazers alters these developmental trajectories (visible as a spread in the data), grazer-induced phenotypes remain constrained within the morphological space of each genotype (ellipses around the phenotypes per genotype/species).

**Table 2.**
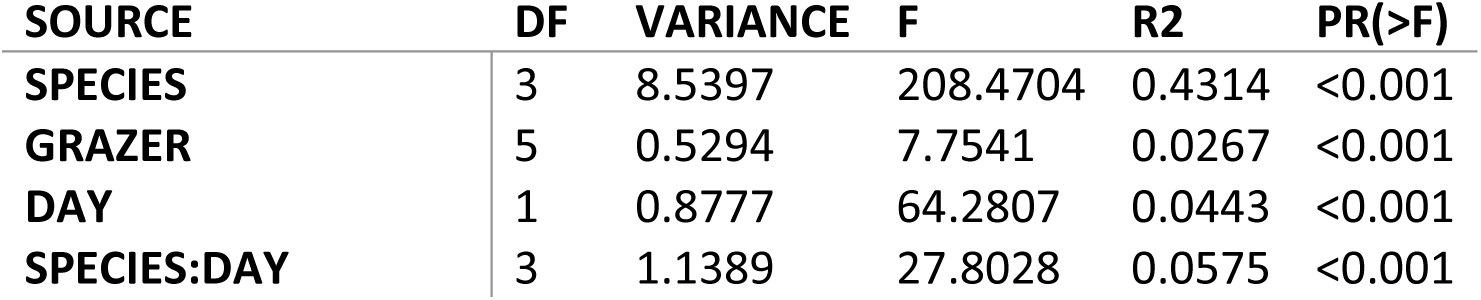
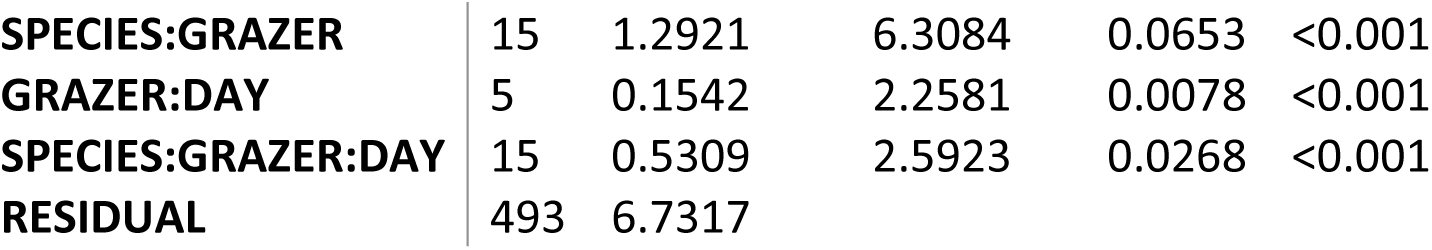
PERMANOVA results with effect sizes for the first PERMANOVA model that includes fungal species, grazer identity and developmental time. The variation in the initial condition for each colony was corrected using PC1 and PC2 co-ordinates for day 1 as co-variates.

| SOURCE | DF | VARIANCE | F | R2 | PR(>F) |
| --- | --- | --- | --- | --- | --- |
| **SPECIES** | 3 | 8.5397 | 208.4704 | 0.4314 | <0.001 |
| **GRAZER** | 5 | 0.5294 | 7.7541 | 0.0267 | <0.001 |
| **DAY** | 1 | 0.8777 | 64.2807 | 0.0443 | <0.001 |
| **SPECIES:DAY** | 3 | 1.1389 | 27.8028 | 0.0575 | <0.001 |
| **SPECIES:GRAZER** | 15 | 1.2921 | 6.3084 | 0.0653 | <0.001 |
| **GRAZER:DAY** | 5 | 0.1542 | 2.2581 | 0.0078 | <0.001 |
| **SPECIES:GRAZER:DAY** | 15 | 0.5309 | 2.5923 | 0.0268 | <0.001 |
| **RESIDUAL** | 493 | 6.7317 |  |  |  |

In the second set of models that focus on the effect of each grazer separately, we did not find support for convergence to a common phenotype. Instead, we found the opposite, that is, we found a significant effect for increasing dispersion among genotypes in grazer-induced morphology (except for Collembola for which the effect was not significant) (Fig. 4). Effect size varied from 5 to 10% and was strongest for woodlice 1 (Oniscus), followed by millipede, woodlice 2 (Porcellio) and nematode treatment. Taken together, these results show first that grazer-induced plasticity was detectable across fungal species, but it was constrained within the trajectory of each genotype and second, that instead of converging to a common phenotype it increased the dissimilarity of the phenotypes.

**Figure 4.**
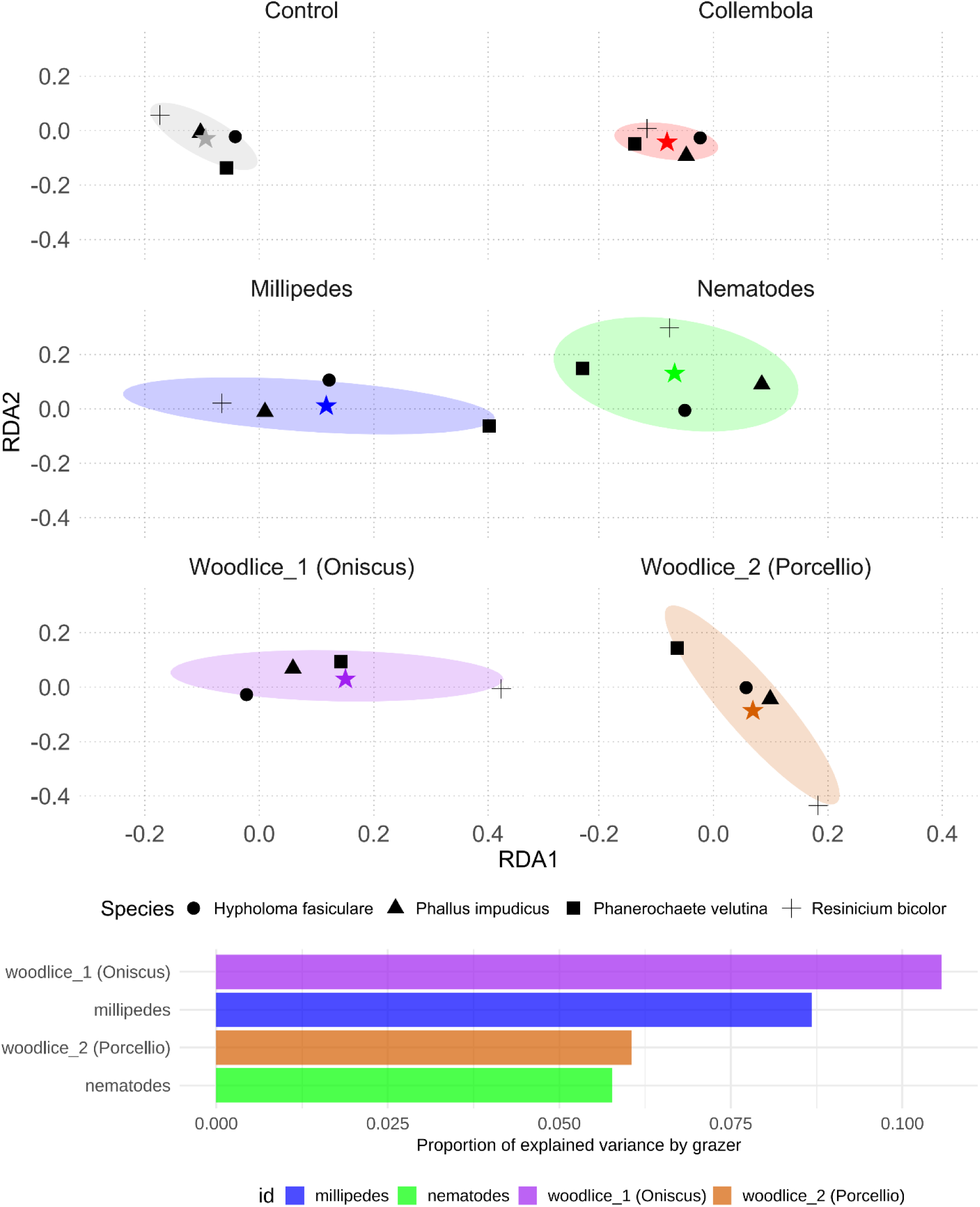
Testing for convergence or divergence in fungal phenotypes under grazer pressure. Each ordination plot shows the average mycelial morphotype of each genotype per grazing treatment (circle = *H. fasciculare*, triangle = *P. impudicus*, square = *P. velutina*, cross = *R. bicolor*). The across-genotype average per grazer treatment is shown as a star. Dispersion around this centroid is visualized with shaded ellipses. With the exception of collembola, where no significant effect was detected, all grazer treatments increased morphological dispersion. Bar plots at the bottom indicate the effect size of grazer-induced changes in dispersion for significant treatments.

### Trajectories of grazer-induced plasticity differed in magnitude and direction among fungal species

We found large variation in the level of plasticity among the fungal species tested across the grazers (Fig. 5). *R. bicolor* exhibited the highest morphological plasticity in response to grazers, with up to 30% of the variation in morphology attributable to grazing (p < 0.01), particularly in response to the two woodlice species (Figs. 6). In both cases, RDA plots revealed clear separation between grazed and ungrazed genotypes of this species (Fig. S2). Multiple traits contributed to this divergence—indicating a broadly distributed response rather than dominance by a single or few traits (Fig. 6). Among the two woodlice, woodlouse 1 (*Oniscus*) had the stronger effect. The overall colony shape remained stable when exposed to this grazer, but the cord morphology shifted to shorter, thinner cords with straighter branching angles. A more prominent change was the increase in cord density, although loop density was unaffected. Despite these structural changes, the predicted transport dynamics remained stable, and robustness even improved in this treatment, particularly against random attacks ortargeted attacks on main cords.

**Figure 5.**
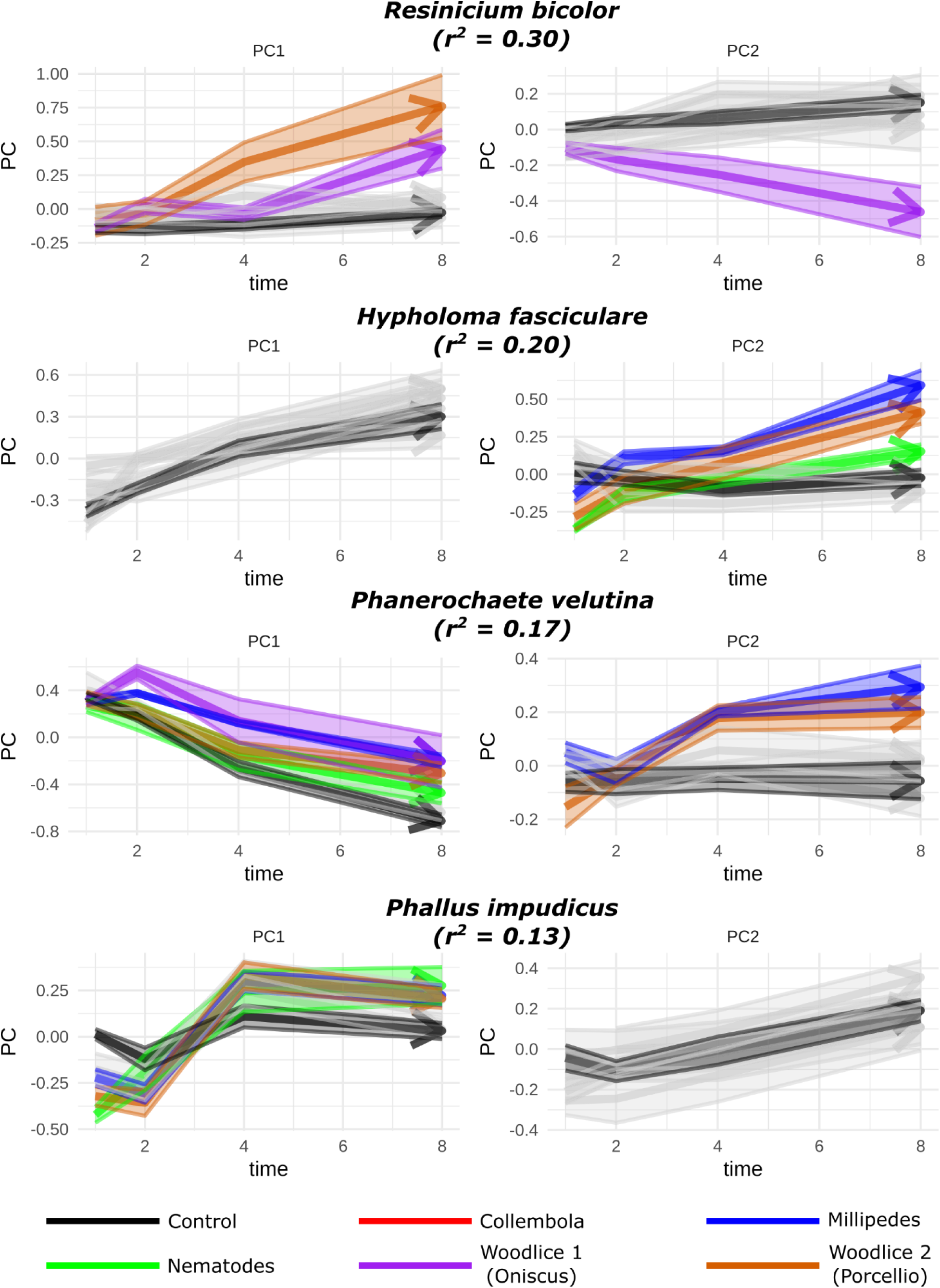
Species-specific trajectories of mycelial morphological development over time and deviations caused by grazer presence. The greatest grazer-induced deviation was observed in *R. bicolor*, where grazers explained 30% of the variation in morphology through time, followed by *H. fasciculare*, *P. velutina*, and *P. impudicus*. Trajectories of grazer-induced morphologies that differ by more than twice the joint standard error from the non-grazed phenotypes are color-coded to highlight these contrasts.

**Figure 6.**
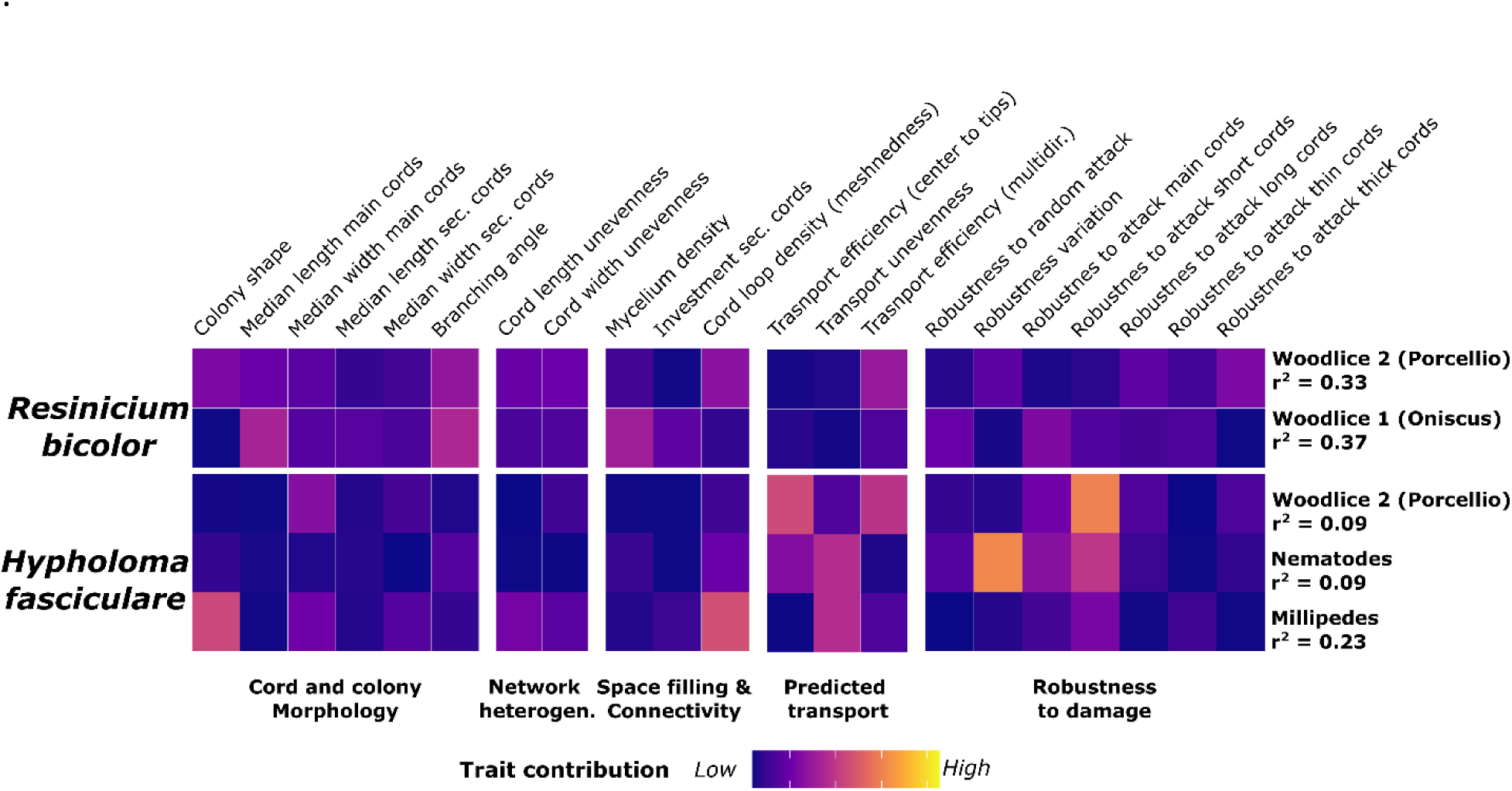
Heatmap of morphological trait shifts between grazed and non-grazed genotypes at the end of the experiment for *R. bicolor* and *H. fasciculare*. These two species showed the strongest grazer-induced deviations in morphology (see Figure 5). For each species, the grazer treatment with the largest deviation is shown, along with the percentage of explained variation. Corresponding RDA biplots can be found in Figure S2.

Colonies exposed to Woodlice 2 (*Porcellio*) became more circular and exhibited similar changes in cord morphology—short, thin cords with straight branching angles. Cord heterogeneity also shifted: cord lengths became more even, while width distribution became more uneven, suggesting the emergence of a few dominant thick cords. In this treatment, the number of loops decreased while density remained stable. These changes were associated with reduced predicted transport efficiency, especially for multidirectional transport, and increased variability in robustness to random attacks. Importantly, robustness to attacks on thick cords declined, indicating these few thick cords were critical for network connectivity and vulnerable to disruption.

*H. fasiculare* showed the second-highest response with ca. 20% (p < 0.05) of morphological variation explained by the presence of the grazer (Fig. 5). This effect was primarily driven by exposure to millipedes with up to 23% explained variation attributed to this grazer. In this case, only a subset of traits shifted, including the development of a more irregular colony shape and a marked decrease in loop number. These structural changes were associated with a reduction in predicted transport efficiency and a sharp increase in the heterogeneity of main transport cords. However, predicted robustness remained largely unaffected, with only a slight increase in robustness to attacks on thin cords. Responses to Woodlice 2 (*Porcellio*) and nematodes were weaker (with around 9% of explained variation attributed to the presence of these grazers (Fig. 6)) and more variable across replicates, as reflected in less consistent RDA separation (Fig. S2).

*P. velutinae* and *P. impudicus* exhibited the lowest morphological plasticity, with only 17% and 13% of trait variation attributed to grazer effects, respectively Fig. 5. Although these effects were statistically significant, RDA plots showed inconsistent separation between treatment groups (Fig. S2), suggesting limited and variable responses across replicates.

## Discussion

We tested whether grazing pressure induced convergence towards a common mycelial phenotype or, alternatively, whether plasticity remained constrained within the trajectories of non-grazed genotypes. We found support for the latter, where despite grazer induced effects altering the morphology of each species, the effect was small and genotypes retained characteristic patterns across all treatments. In fact, we found that changes induced led to increased dissimilarities among grazed genotypes rather than convergence to a particular grazing-resistant phenotype, as observed for grazing in other kingdoms. These results are surprising based on the assumption that fungi have the capacity to vary their morphology significantly (extreme plasticity) in comparison to that of animals and plants (Slepecky & Starmer 2009). These results suggest that the mycelium morphology has a more limited flexibility to reconfigure itself than previously thought. It also suggests that morphological plasticity may not be the most effective means to deal with grazing pressure.

### Strong species differences suggest limits in morphological plasticity

Although some of our species exhibited considerable plasticity in response to grazers, these changes were not strong enough to override genotype-specific differences. In other words, the morphological trajectories of each fungal genotype remained constrained within distinct boundaries defined by genotype identity. This pattern suggests that the plasticity of cord-forming fungi is constrained within certain species-specific limits, at least in response to grazing. In addition, these limits varied among species, with *R. bicolor* showing up to 30% variation in morphology due to grazing pressure, while *P. impudicus* exhibited only around 10% change in response to grazers.

The low levels of plasiticity in *P. impudicus* might be explained by several factors. It could be that the architecture of *P. impudicus* constitutively embodies grazing-resistant features, such as a high level of cross-linking, however this explanation is unlikely as there was no conversion into a common grazed phenotype for the other species. In theory, when genotypes show low levels of grazer-induced plasticity it can explained by the lack of sensory systems to detect and respond to grazers (Murren *et al*. 2015) or show variation in the metabolic cost of changing the phenotype (Auld *et al*. 2010). While we do not have direct evidence of sensory mechanisms for our species, we expect they are present as they are in closely related taxa (e.g. *Coprionopsis cinerea* and *Pleurotus ostreatus*) are known to respond to at least nematodes and collembola (Plaza *et al*. 2016; Schmieder *et al*. 2019). Thus, more likely, lower levels of plasticity may be associated with variation in the metabolic cost of altering the morphology of mycelial phenotype sufficiently to achieve resistance to grazing in comparison to allocation to other forms of defence. We speculate these costs may reflect differences in the construction of cords. There is evidence that the level complexity in the histology of the cords varies across species (Thompson & Rayner 1983; Townsend 1954). For example, *P. impudicus,* the least plastic species in our study, is at the more complex end of the spectrum with clear differentiation of anatomical regions within the cord compared to that of other cord-forming fungi (Eamus *et al*. 1985; Townsend 1954). Likewise, *P. velutina*, which also shows relatively low plasticity in this study, has also been reported as showing zone differentiation in its cord anatomy (Thompson & Rayner 1983). In contrast *H. fasciculare*, with relative higher levels of plasticity in our study, is reported to on the lower end of cord complexity (Thompson & Rayner 1983). We have not found anatomical studies of the cords of *R. bicolor* (the species with the most plasticity in our study), but it is a fast-growing-species (Zakaria & Boddy 2002) a life-history trait that is commonly associated with lower investment in resources in the construction of an organism’s body (Wright *et al*. 2004). Thus, we hypothesized with increasing complexity of the cord-forming mycelia is part of a trade-off with the ability to respond to grazer pressure. For species with less plastic and more complex phenotypes, modifying the network architecture may jeopardize efficient transport which might outweigh any benefit to morphological responses to grazing.

We also emphasize that these observed limits to grazer-induced plasticity may not reflect the potential for morphological plasticity in response to other environmental factors. For example, we observed relatively high variation of mycelium morphology at the start of the experiment within replicates of the same genotype reflecting initial differences in mycelial growth before the addition of grazers (Fig. S3). Such differences may arise from inherent heterogeneity in resource quality, or the extent colonization of wood blocks used as inoculum in these experiments. Such differences may have long lasting effect determining the trajectory of the developing mycelia. Such effects have been reported for cord-forming fungi, where the intensity of colonization of wood blocks can strongly the development of the mycelium even across generations (Fukasawa *et al*. 2020). Overall, while our results indicate limits to grazer-induced plasticity in cord-forming fungi, they also suggest that other factors such as resource heterogeneity may drive greater levels of morphological plasticity.

### Lack of convergence may be due to weak evolutionary pressure of fungal grazing for induced morphological plasticity

Although we cannot determine whether the observed grazer-induced plasticity is adaptive— such a study would require long-term monitoring of fitness consequences of plastic phenotypes—our findings do not support that fungal grazing exerts strong evolutionary pressure for the development of adaptive morphology. We found no evidence of a convergent ’grazing-resistant phenotype’ across species and instead observed substantial variation in plastic responses. The emergence of a common adaptive phenotype typically requires strong and consistent environmental pressures. Given the diverse community of fungal grazers present in soils, optimizing morphology for one grazer may offer limited fitness benefits if it compromises adaptability to other grazers or environmental factors (Einum *et al*. 2019). Even if such optimal phenotype exist, the fact that we did not observe a response in our time window indicates that individuals would spend a significant part of their life cycle in a sub-optimal transitioning phenotype, a feature of weak selection pressure (Einum & Burton 2023).

We also note, however, that grazing can still impose strong evolutionary pressure for the evolution of plasticity in other traits that are not morphological. Phenotypic convergence could be exhibited in traits not related to morphology that we did not measure in this study as “hidden plasticity” (Forsman 2015), such as production of chemical compounds that reduce palatability and intoxicate the grazers. This type of responses is well documented in in plants (Fernández De Bobadilla *et al*. 2022) and more recently it has been found that some fungi produce compounds with similar functionality (Lee *et al*. 2020; Plaza *et al*. 2016; Schmieder *et al*. 2019). So, while the apparent target morphological phenotype between grazed and ungrazed environment are different, “hidden” grazer induced plasticity in the form of altered chemistry may act as compensatory mechanism to maintain the apparent dissimilar phenotype between the two environments (Metcalfe 2024).

### Implications of plasticity limits in trait-based fungal ecology

The significant differences between species but limited morphological plasticity with species suggest that mycelium morphology embodies useful functional traits to understand community assembly and ecosystem processes (Mcgill *et al*. 2006). In community ecology, traits that exhibit lower within-species than among-species variation can reflect consistent ecological strategies and predict patterns of species composition along environmental gradients (Franklin *et al*. 2020; HilleRisLambers *et al*. 2012). For example, lab-derived traits in marine plankton and zooplankton have successfully predicted seasonal shifts and habitat filtering in natural settings (Edwards *et al*. 2013; Vogt *et al*. 2013). If fungal mycelial traits reflect species differences, they could also predict how shifts in fungal community composition along environmental gradients impact ecosystem processes, such as carbon cycling. Cord-forming fungi, for instance, play a key role in lignin decomposition, and morphological traits may influence decomposition efficiency, thereby affecting carbon budgets. Although speculative, these findings suggest that predictable relationships between mycelial morphology, environmental conditions, and ecosystem functions warrant further exploration.

### Implications of using fungi as model systems for studying the ecology and evolution of plasticity

Moving forward, it remains to be tested whether non-cord-forming fungi exhibit the expected extreme plasticity attributed to fungi (Klein & Paschke 2004; Slepecky & Starmer 2009). Given that phenotypic plasticity tends to evolve in highly variable environments (Lande 2009), fungi inhabiting soils—among the most heterogeneous habitats—would be expected to show strong selective pressures for high morphological plasticity. Evolutionary models suggest that filamentous growth forms, which are particularly advantageous in spatially heterogeneous environments like soils, would not have evolved in more homogeneous aquatic habitats (Heaton *et al*. 2020). This parallels findings in ectotherms, where greater plasticity is observed in terrestrial species compared to aquatic ones, likely due to higher environmental variability on land (Einum & Burton 2023). However, our results suggest limited plasticity in cord-forming fungi, a phenotype that has evolved repeatedly in the Basidiomycota (Monk 2014). One possible explanation for this reduced plasticity is that cord-forming fungi, with their complex and energetically costly growth structures, may have evolved to optimize efficiency in specific ecological roles. This specialization could limit the need for extensive morphological plasticity. Understanding the evolutionary pressures that led to the emergence and maintenance of such behavior in less plastic fungi would be crucial, particularly in relation to their role in ecosystem processes like carbon cycling.

Overall, our study highlights how fungi can expand our understanding of plasticity. Existing theories, largely developed for organisms with deterministic body plans, need refinement to better explain how phenotypic plasticity emerges and is maintained in organisms with complex, adaptable structures that evolved in variable environments. Our findings suggest that investigating these questions in cord-forming fungi may provide new insights into the constraints of morphological plasticity under diverse and dynamic ecological conditions.

## Conflict of interest

All authors declare no conflict of interest.

## Disclosure

CAAT are funded by an Academy Research Fellowship from the Research Council of Finland (RCF 356191) and a Feodor-Lynnen Fellowship by the Alexander von Humboldt Foundation. MDF acknowledges funding from the Leverhulme Trust (RPG-2015-437) and the OxBer partnership fund.

## Author contributions

CAAT and MDF conceived the paper. LB provided images of experiments. MDF performed the image analysis, CAAT did the statistical analysis. CAAT wrote the first version of the manuscript. All coauthors contributed in writing the final version of the manuscript.

## Acknowledgements

We thank undergraduate students Jonathan Heale, Geoffrey Liddell, Elizabeth McGowan and Freya Way at the University of Oxford for help in the development of the image processing pipeline.

## Notes

### Competing Interest Statement

The authors have declared no competing interest.

### Summary of Updates

Conceptual and data figures were updates, authors affiliations updated and discusion was updated to clarify observed differences in plasticity.

## References

Adams, D.C. & Collyer, M.L. (2009). A GENERAL FRAMEWORK FOR THE ANALYSIS OF PHENOTYPIC TRAJECTORIES IN EVOLUTIONARY STUDIES. Evolution, 63, 1143–1154.

Aguilar-Trigueros, C.A., Boddy, L., Rillig, M.C. & Fricker, M.D. (2022). Network traits predict ecological strategies in fungi. ISME Communications, 2, 2.

Alster, C.J., Allison, S.D., Johnson, N.G., Glassman, S.I. & Treseder, K.K. (2021). Phenotypic plasticity of fungal traits in response to moisture and temperature. ISME Communications, 1, 43.

Anthony, M.A., Bender, S.F. & Van Der Heijden, M.G.A. (2023). Enumerating soil biodiversity. Proc. Natl. Acad. Sci. U.S.A., 120, e2304663120.

Auld, J.R., Agrawal, A.A. & Relyea, R.A. (2010). Re-evaluating the costs and limits of adaptive phenotypic plasticity. Proc. R. Soc. B., 277, 503–511.

Auld, J.R. & Relyea, R.A. (2011). Adaptive plasticity in predator-induced defenses in a common freshwater snail: altered selection and mode of predation due to prey phenotype. Evol Ecol, 25, 189–202.

Behm, J.E. & Kiers, E.T. (2014). A phenotypic plasticity framework for assessing intraspecific variation in arbuscular mycorrhizal fungal traits. Journal of Ecology, 102, 315–327.

Bielčik, M., Aguilar-Trigueros, C.A., Lakovic, M., Jeltsch, F. & Rillig, M.C. (2019). The role of active movement in fungal ecology and community assembly. Mov Ecol, 7, 36.

Boddy, L. (1999). Saprotrophic cord-forming fungi: meeting the challenge of heterogeneous environments. Mycologia, 91, 13–32.

Boyce, K.J. & Andrianopoulos, A. (2015). Fungal dimorphism: the switch from hyphae to yeast is a specialized morphogenetic adaptation allowing colonization of a host. FEMS Microbiology Reviews, 39, 797–811.

Coleine, C., Stajich, J.E. & Selbmann, L. (2022). Fungi are key players in extreme ecosystems. Trends in Ecology & Evolution, 37, 517–528.

Crowther, T.W., Boddy, L. & Hefin Jones, T. (2012). Functional and ecological consequences of saprotrophic fungus–grazer interactions. The ISME Journal, 6, 1992–2001.

Crowther, T.W., Boddy, L. & Jones, T.H. (2011). Species-specific effects of soil fauna on fungal foraging and decomposition. Oecologia, 167, 535–545.

Csardi G & Nepusz T. (2006). The igraph software package for complex network research. *InterJournal*, Complex Systems, 1695.

Dikec, J., Olivier, A., Bobée, C., D’Angelo, Y., Catellier, R., David, P., et al. (2020). Hyphal network whole field imaging allows for accurate estimation of anastomosis rates and branching dynamics of the filamentous fungus Podospora anserina. Sci Rep, 10, 3131.

Dorey, T. & Schiestl, F.P. (2022). Plant phenotypic plasticity changes pollinator-mediated selection. Evolution, evo.14634.

Du, H., Lv, P., Ayouz, M., Besserer, A. & Perré, P. (2016). Morphological Characterization and Quantification of the Mycelial Growth of the Brown-Rot Fungus Postia placenta for Modeling Purposes. PLOS ONE, 11, e0162469.

Eamus, D., Thompson, W., Cairney, J.W.G. & Jennings, D.H. (1985). Internal Structure and Hydraulic Conductivity of Basidiomycete Translocating Organs. J Exp Bot, 36, 1110–1116.

Edwards, K.F., Litchman, E. & Klausmeier, C.A. (2013). Functional traits explain phytoplankton community structure and seasonal dynamics in a marine ecosystem. Ecology Letters, 16, 56–63.

Einum, S. & Burton, T. (2023). Divergence in rates of phenotypic plasticity among ectotherms. Ecology Letters, 26, 147–156.

Einum, S., Ratikainen, I., Wright, J., Pélabon, C., Bech, C., Jutfelt, F., et al. (2019). How to quantify thermal acclimation capacity? Global Change Biology, 25, 1893–1894.

Fernández De Bobadilla, M., Vitiello, A., Erb, M. & Poelman, E.H. (2022). Plant defense strategies against attack by multiple herbivores. Trends in Plant Science, 27, 528–535.

Flenner, I., Olne, K., Suhling, F. & Sahlén, G. (2009). Predator-induced spine length and exocuticle thickness in *Leucorrhinia dubia* (Insecta: Odonata): a simple physiological trade-off? Ecological Entomology, 34, 735–740.

Forsman, A. (2015). Rethinking phenotypic plasticity and its consequences for individuals, populations and species. Heredity, 115, 276–284.

Franklin, O., Harrison, S.P., Dewar, R., Farrior, C.E., Brännström, Å., Dieckmann, U., et al. (2020). Organizing principles for vegetation dynamics. Nat. Plants, 6, 444–453.

Fricker, Boddy, L. & Bebber, D. (2007). Network Organisation of Mycelial Fungi. In: Biology of the Fungal Cell, The Mycota (eds. Howard, R.J. & Gow, N.A.R.). Springer Berlin Heidelberg, Berlin, Heidelberg, pp. 309–330.

Fricker, Heaton, Luke L. M, Jones, Nick S., & Boddy, Lynne. (2017a). The Mycelium as a Network. Microbiology Spectrum, 5, 10.1128/microbiolspec.funk-0033–2017.

Fricker, M.D., Akita, Heaton, L.L.M., Jones, N., Obara, B. & Nakagaki, T. (2017b). Automated analysis of Physarum network structure and dynamics. Journal of Physics D: Applied Physics, 50, 254005.

Fukasawa, Y., Savoury, M. & Boddy, L. (2020). Ecological memory and relocation decisions in fungal mycelial networks: responses to quantity and location of new resources. The ISME Journal, 14, 380–388.

Gauthier, G.M. (2015). Dimorphism in Fungal Pathogens of Mammals, Plants, and Insects. PLoS Pathog, 11, e1004608.

Gilbert, J.J. (2013). The cost of predator-induced morphological defense in rotifers: experimental studies and synthesis. Journal of Plankton Research, 35, 461–472.

Gomulkiewicz, R. & Stinchcombe, J.R. (2022). Phenotypic plasticity made simple, but not too simple. American J of Botany, 109, 1519–1524.

Heaton, L.L.M., Jones, N.S. & Fricker, M.D. (2020). A mechanistic explanation of the transition to simple multicellularity in fungi. Nat Commun, 11, 2594.

Heaton, L.L.M., López, E., Maini, P.K., Fricker, M.D. & Jones, N.S. (2012a). Advection, diffusion, and delivery over a network. Phys. Rev. E, 86, 021905.

Heaton, L.L.M., López, Eduardo., Fricker, M.D. & Jones, N.S. (2010). Growth-induced mass flows in fungal networks. Proceedings of the Royal Society B: Biological Sciences, 277, 3265– 3274.

Heaton, L.L.M., Obara, B., Grau, V., Jones, N., Nakagaki, T., Boddy, L., et al. (2012b). Analysis of fungal networks. Fungal Biology Reviews, 26, 12–29.

HilleRisLambers, J., Adler, P.B., Harpole, W.S., Levine, J.M. & Mayfield, M.M. (2012). Rethinking Community Assembly through the Lens of Coexistence Theory. Annu. Rev. Ecol. Evol. Syst., 43, 227–248.

Hoverman, J.T. & Relyea, R.A. (2007). HOW FLEXIBLE IS PHENOTYPIC PLASTICITY? DEVELOPMENTAL WINDOWS FOR TRAIT INDUCTION AND REVERSAL. Ecology, 88, 693–705.

Klein, D.A. & Paschke, M.W. (2004). Filamentous Fungi: the Indeterminate Lifestyle and Microbial Ecology. Microb Ecol, 47.

Lande, R. (2009). Adaptation to an extraordinary environment by evolution of phenotypic plasticity and genetic assimilation. J of Evolutionary Biology, 22, 1435–1446.

Lebbink, G., Risch, A.C., Schuetz, M. & Firn, J. (2024). How plant traits respond to and affect vertebrate and invertebrate herbivores—Are measurements comparable across herbivore types? Plant Cell & Environment, 47, 5–23.

Lee, C.-H., Chang, H.-W., Yang, C.-T., Wali, N., Shie, J.-J. & Hsueh, Y.-P. (2020). Sensory cilia as the Achilles heel of nematodes when attacked by carnivorous mushrooms. Proc. Natl. Acad. Sci. U.S.A., 117, 6014–6022.

Lopez-Molina, C., Vidal-Diez De Ulzurrun, G., Baetens, J.M., Van Den Bulcke, J. & De Baets, B. (2015). Unsupervised ridge detection using second order anisotropic Gaussian kernels. Signal Processing, 116, 55–67.

Mcgill, B., Enquist, B., Weiher, E. & Westoby, M. (2006). Rebuilding community ecology from functional traits. Trends in Ecology & Evolution, 21, 178–185.

Metcalfe, N.B. (2024). How important is hidden phenotypic plasticity arising from alternative but converging developmental trajectories, and what limits it? Journal of Experimental Biology, 227, jeb246010.

Monk, K.K. (2014). INVESTIGATING THE ECOLOGY, DIVERSITY AND DISTRIBUTION OF CORD-FORMING FUNGI IN GREAT BRITAIN. Linacre College, University of Oxford.

Murren, C.J., Auld, J.R., Callahan, H., Ghalambor, C.K., Handelsman, C.A., Heskel, M.A., et al. (2015). Constraints on the evolution of phenotypic plasticity: limits and costs of phenotype and plasticity. Heredity, 115, 293–301.

Obara, B., Grau, V. & Fricker, M.D. (2012). A bioimage informatics approach to automatically extract complex fungal networks. Bioinformatics, 28, 2374–2381.

Oksanen, J., Kindt, R., Legendre, P., O’Hara, B., Stevens, M.H.H., Oksanen, M.J., et al. (2007). The vegan package. Community ecology package, 10, 719.

Oyarte Galvez, L., Bisot, C., Bourrianne, P., Cargill, R., Klein, M., van Son, M., et al. (2025). A travelling-wave strategy for plant–fungal trade. Nature.

Pain, C., Kriechbaumer, V., Kittelmann, M., Hawes, C. & Fricker, M. (2019). Quantitative analysis of plant ER architecture and dynamics. Nature Communications, 10, 984.

Pfennig, D.W. (2021). Phenotypic Plasticity & Evolution: Causes, Consequences, Controversies. 1st edn. CRC Press, Boca Raton.

Pigliucci, M. & Müller, G.B. (Eds.). (2010). Evolution, the Extended Synthesis. The MIT Press.

Plaza, D.F., Schmieder, S.S., Lipzen, A., Lindquist, E. & Künzler, M. (2016). Identification of a Novel Nematotoxic Protein by Challenging the Model Mushroom *Coprinopsis cinerea* with a Fungivorous Nematode. G3 Genes|Genomes|Genetics, 6, 87–98.

R Core Team, R. (2013). R: A language and environment for statistical computing.

Rotheray, T.D., Jones, T.H., Fricker, M.D. & Boddy, L. (2008). Grazing alters network architecture during interspecific mycelial interactions. Fungal Ecology, 1, 124–132.

Schmieder, S.S., Stanley, C.E., Rzepiela, A., Van Swaay, D., Sabotič, J., Nørrelykke, S.F., et al. (2019). Bidirectional Propagation of Signals and Nutrients in Fungal Networks via Specialized Hyphae. Current Biology, 29, 217–228.e4.

Schneider, H.M. (2022). Characterization, costs, cues and future perspectives of phenotypic plasticity. Annals of Botany, 130, 131–148.

Sentis, A., Bertram, R., Dardenne, N., Ramon-Portugal, F., Louit, I., Le Trionnaire, G., et al. (2019). Different phenotypic plastic responses to predators observed among aphid lineages specialized on different host plants. Sci Rep, 9, 9017.

Slepecky, R.A. & Starmer, W.T. (2009). Phenotypic plasticity in fungi: a review with observations on *Aureobasidium pullulans*. Mycologia, 101, 823–832.

Spitze, K. & Sadler, T.D. (1996). Evolution of a Generalist Genotype: Multivariate Analysis of the Adaptiveness of Phenotypic Plasticity. The American Naturalist, 148, S108–S123.

Sten, O., Del Dottore, E., Pugno, N. & Mazzolai, B. (2024). A ridge-based detection algorithm with filament overlap identification for 2D mycelium network analysis. Ecological Informatics, 82, 102670.

Sultan, S.E. (2003). Phenotypic plasticity in plants: a case study in ecological development. Evolution and Development, 5, 25–33.

Sultan, S.E. (2021). Phenotypic plasticity as an intrinsic property of organisms. In: Phenotypic plasticity & evolution. CRC Press, pp. 3–24.

Thompson, W. & Rayner, A.D.M. (1983). Extent, development and function of mycelial cord systems in soil. Transactions of the British Mycological Society, 81, 333–345.

Tlalka, M., Watkinson, S.C., Darrah, P.R. & Fricker, M.D. (2002). Continuous imaging of amino-acid translocation in intact mycelia of *Phanerochaete velutina* reveals rapid, pulsatile fluxes. New Phytologist, 153, 173–184.

Tollrian, R. & Harvell, C.D. (1999). The ecology and evolution of inducible defenses. Princeton university press.

Tordoff, G.M., Boddy, L. & Jones, T.H. (2006). Grazing by Folsomia candida (Collembola) differentially affects mycelial morphology of the cord-forming basidiomycetes Hypholoma fasciculare, Phanerochaete velutina and Resinicium bicolor. Mycological Research, 110, 335–345.

Tordoff, G.M., Boddy, L. & Jones, T.H. (2008). Species-specific impacts of collembola grazing on fungal foraging ecology. Soil Biology and Biochemistry, 40, 434–442.

Townsend, B.B. (1954). Morphology and development of fungal Rhizomorphs. Transactions of the British Mycological Society, 37, 222–233.

Ugalde, U. & Rodriguez-Urra, A.B. (2014). The Mycelium Blueprint: insights into the cues that shape the filamentous fungal colony. Appl Microbiol Biotechnol, 98, 8809–8819.

Veresoglou, S.D., Wang, D., Andrade-Linares, D.R., Hempel, S. & Rillig, M.C. (2018). Fungal Decision to Exploit or Explore Depends on Growth Rate. Microb Ecol, 75, 289–292.

Vidal-Diez De Ulzurrun, G., Baetens, J.M., Van Den Bulcke, J., Lopez-Molina, C., De Windt, I. & De Baets, B. (2015). Automated image-based analysis of spatio-temporal fungal dynamics. Fungal Genetics and Biology, 84, 12–25.

Vogt, R.J., Peres-Neto, P.R. & Beisner, B.E. (2013). Using functional traits to investigate the determinants of crustacean zooplankton community structure. Oikos, 122, 1700–1709.

West-Eberhard, M.J. (1989). Phenotypic Plasticity and the Origins of Diversity. Annual Review of Ecology and Systematics, 20, 249–278.

Wickham, H., Averick, M., Bryan, J., Chang, W., McGowan, L., François, R., et al. (2019). Welcome to the Tidyverse. JOSS, 4, 1686.

Wright, I.J., Reich, P.B., Westoby, M., Ackerly, D.D., Baruch, Z., Bongers, F., et al. (2004). The worldwide leaf economics spectrum. Nature, 428, 821–827.

Xu, H., Blonder, B., Jodra, M., Malhi, Y. & Fricker, M. (2021). Automated and accurate segmentation of leaf venation networks via deep learning. New Phytologist, 229, 631– 648.

Yafetto, L. (2018). The structure of mycelial cords and rhizomorphs of fungi: A minireview. Mycosphere, 9, 984–998.

Yin, X., Jin, W., Zhou, Y., Wang, P. & Zhao, W. (2017). Hidden defensive morphology in rotifers: benefits, costs, and fitness consequences. Sci Rep, 7, 4488.

Zakaria, A.J. & Boddy, L. (2002). Mycelial foraging by Resinicium bicolor: interactive effects of resource quantity, quality and soil composition. FEMS Microbiology Ecology, 40, 135– 142.

