## Supplementary material for "Not Extremely Plastic: Testing the Limits of Morphological Plasticity in Fungal Mycelia in Response to Soil Grazers": Table S1; Fig S1, Fig S2, Fig S3

**Table S1. Extended description of the five trait types used in this study**. These traits summarize different patterns of the whole mycelium and are measured through multiple metrics reflecting the construction of the network, density, average and variation in expected transport and robustness to change.

| Type | Biological meaning | Trait | Metric |
|---|---|---|---|
| 1. **Cord and colony morphology** | Size and shape of the whole colony and of the cords | Median cord length and width (main and secondary routes), median branching angle, colony circularity. | (1) Colony circularity, (2) median length of main cords, (3) median length of secondary cords, (4) median width of main cords; (5) median width of secondary cords; (6) branching angle. |
| 2. **Network heterogeneity** | Variation in how fungi build cords within the network | Skewness of cord width and length of main routes. | (7) Skewness length of main cords; (8) skewness width of main cords. |
| 3. **Space filling and connectivity** | Area explored for resources and extent of network interconnection | Mycelial density, loop density (meshnedness), investment in secondary routes. | (9) Cord length per unit of area; (10) alpha coefficient*; (11) ratio total mycelial volume to mycelial volume main cords. |
| 4. **Predicted transport** | Estimated average transport flow through cords and its variation within the network | Expected average transport from inoculum to margin, expected multidirectional transport, skewness of transport from inoculum to margin. | (12) expected transport efficiency to mycelium front; (13) expected transport efficiency through the network; (14) skewness of expected transport efficiency of main cords |
| 5. **Predicted robustness to damage** | Expected robustness to removal of cords of the network | Percentage of cords that can be removed before network volume drops below 50%; removal scenarios include random, targeted by cord width, and targeted by cord length. | (15) robustness to attack of main cords; (16) robustness to random attack of the network; (17) variation in robustness to random attack, robustness to attack of thin (18), thick (19), short (20), and long (21) cords. |

* Here, the alpha coefficient (also referred to as *meshedness*) quantifies the number of loops formed by cords relative to the overall structure of the fungal network. In this context, a *loop* is defined as a closed structure formed by at least three interconnected cords. The alpha coefficient is a metric derived from network science that estimates the density of such loops, calculated using the formula: (ecount(x) – vcount(x) + 1) / (2 × vcount(x) – 5), where *ecount(x)* is the number of cords (edges) and *vcount(x)* is the number of nodes (branching and tip points) in the network.

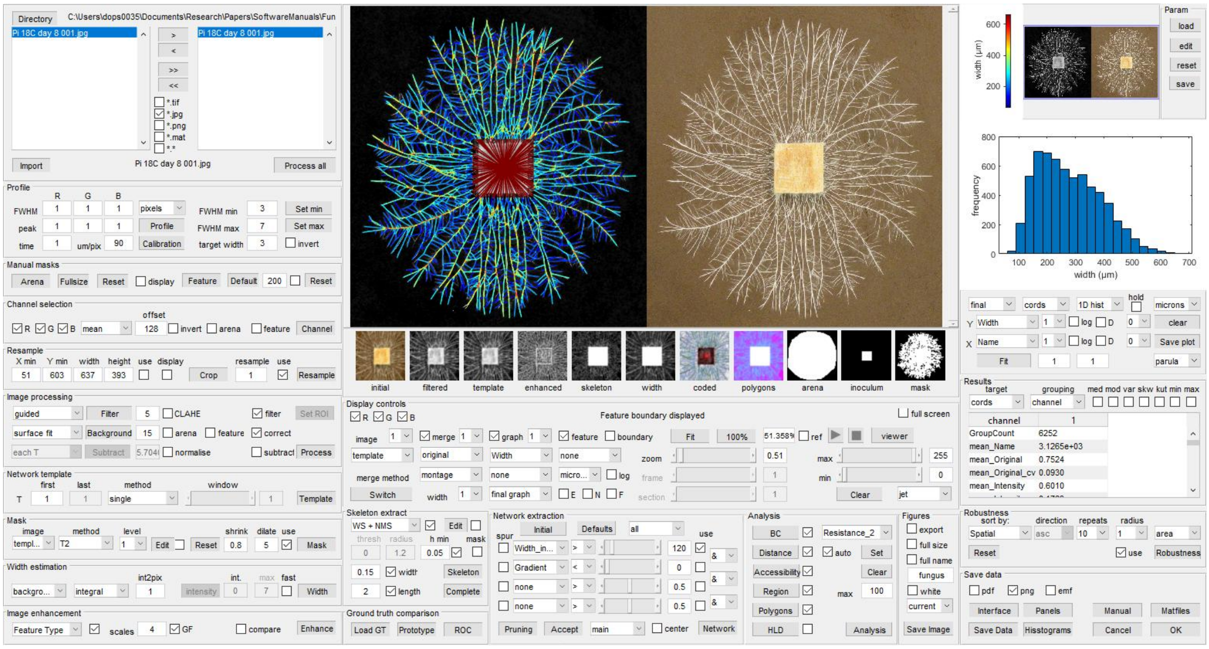

Figure S1. GUI

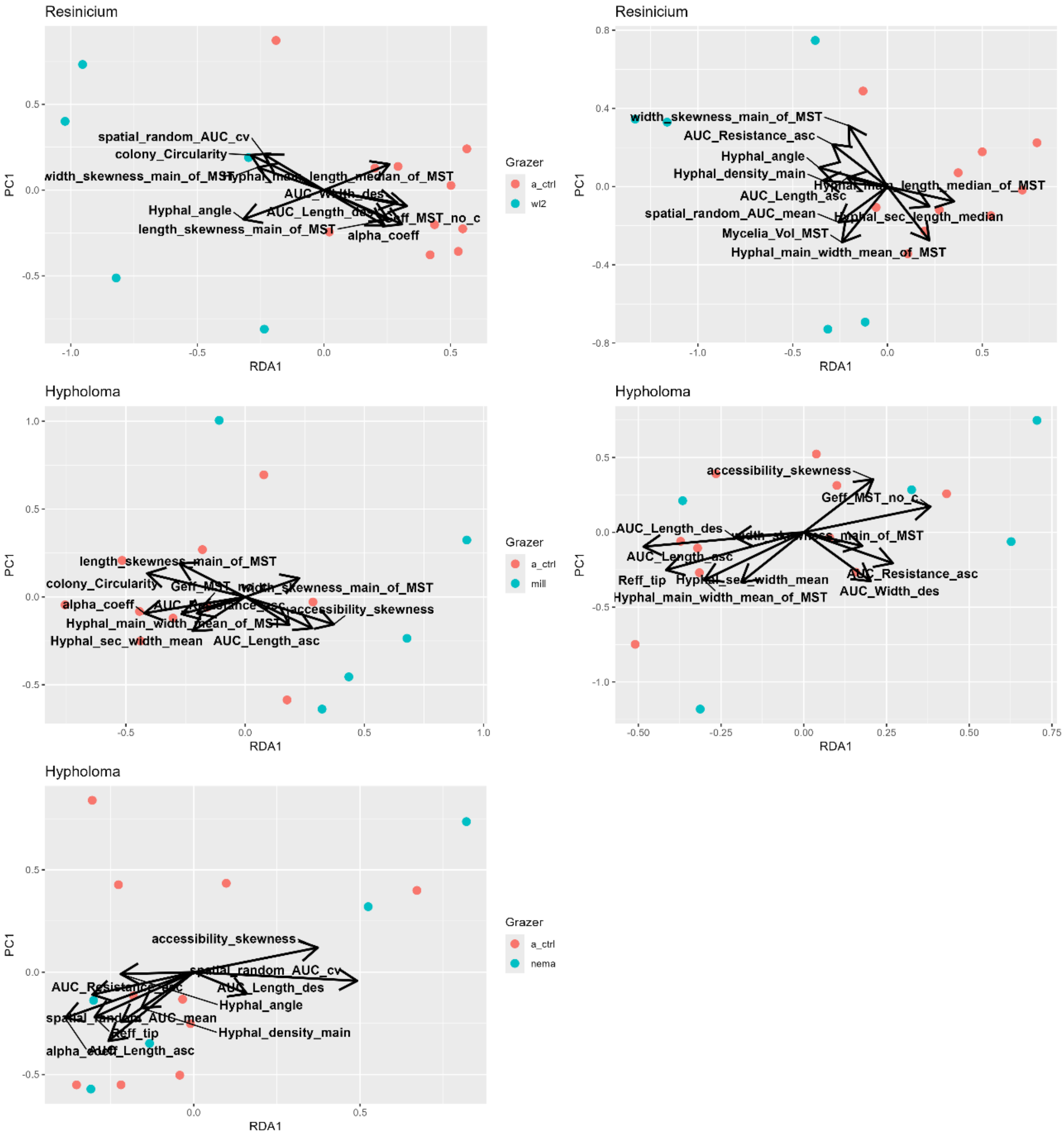


Figure S2. RDA plots showing separation of fungi with highest effect sizes.

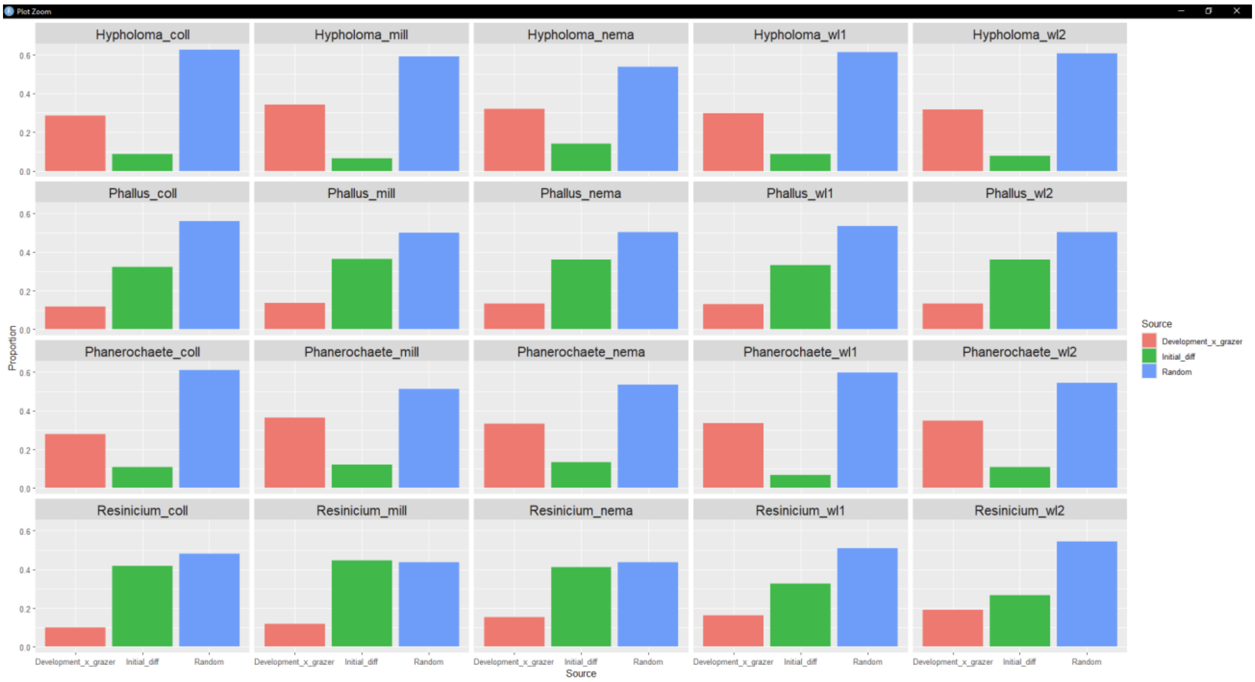


Fig S3. Variance partition among variables in the RDA. Each bar plot represents the proportion of explained variation by the explanatory variables in the RDA (that is, by the time and presence/absence of grazers), the initial growth before the addition of grazer and unaccounted (random) variation.
